# Reduced cortico-accumbal excitatory input due to Nav1.2 haploinsufficiency impairs sociability independently of dopamine

**DOI:** 10.64898/2026.04.15.718826

**Authors:** Toshimitsu Suzuki, Shiori Tominaga, Yuto Yokoi, Hiroaki Mizukami, Kenta Kobayashi, Wakana Nishida, Kona Yamashita, Takahiko Kondo, Yurina Hibi, Tetsushi Yamagata, Shigeyoshi Itohara, Hiroshi Nomura, Hideki Hida, Kazuhiro Yamakawa

## Abstract

Mutations in *SCN2A*, which encodes the voltage-gated sodium channel Nav1.2, are associated with a wide spectrum of neurodevelopmental and neuropsychiatric disorders, including epilepsy, autism spectrum disorder (ASD), and schizophrenia. Although dysfunction of *SCN2A*-dependent neural circuits has been implicated in these disorders, the circuit mechanisms underlying social behavioral abnormalities remain poorly understood. Here, we investigated the neural circuit basis of social behavioral deficits associated with *Scn2a* dysfunction, focusing on the nucleus accumbens (NAc), a key hub in cortico-limbic circuits that regulates emotional and motivational behaviors. Using conditional genetic and chemogenetic approaches in mice, we examined the roles of dorsal telencephalic excitatory neurons, including those in the cerebral cortex, hippocampus, and amygdala, as well as parvalbumin-positive fast-spiking interneurons (PV⁺ FSIs) in the NAc. Mice with *Scn2a* haploinsufficiency in dorsal telencephalic excitatory neurons (*Scn2a*^fl/+^/*Emx1*-Cre) exhibited reduced sociability in the three-chamber social interaction test. Similarly, chemogenetic inhibition of NAc PV⁺ FSIs decreased sociability without affecting locomotor activity or anxiety-like behavior. *Scn2a*^fl/+^/*Emx1*-Cre mice also showed a trend toward reduced prepulse inhibition of the acoustic startle response. Notably, dopamine release into the NAc in the *Scn2a*^fl/+^/*Emx1*-Cre and systemic *Scn2a* heterozygous knockout (*Scn2a*^+/-^) mice was largely comparable to that in control mice. Together, these findings indicate that reduced activity of dorsal telencephalic excitatory neurons or NAc PV⁺ FSIs is sufficient to impair sociability independently of mesolimbic dopamine hypofunction. Our results highlight a potential role of cortico-accumbal circuits in social behavioral deficits associated with *SCN2A* dysfunction.

## Introduction

The voltage-gated sodium channel Nav1.2, encoded by *SCN2A*, plays a critical role in the initiation and propagation of action potentials in excitatory neurons, thereby shaping cortical network excitability and information processing [Catterall, 1992; Catterall *et al*., 2005]. Nav1.2 is abundantly expressed in the axons of cortical and hippocampal excitatory neurons [Liao *et al*., 2010; Ogiwara *et al*., 2018; Yamagata *et al*., 2023], as well as in dopaminergic neurons in the substantia nigra pars compacta and ventral tegmental area (VTA) [Yang *et al*., 2019]. In addition, Nav1.2 is present in specific populations of inhibitory neurons, including medium spiny neurons (MSNs) in the striatum [Miyazaki *et al*., 2014] and GABAergic interneurons in the neocortex [Li *et al*., 2014; Yamagata *et al*., 2017].

Pathogenic mutations in *SCN2A* have been implicated in a broad spectrum of neurodevelopmental and neuropsychiatric disorders, including epilepsy, autism spectrum disorder (ASD), intellectual disability (ID), and schizophrenia [Sugawara *et al*, 2001; Kamiya *et al*, 2004; Ogiwara *et al*, 2009; Buxbaum *et al*, 2012; Rauch *et al*, 2012; de Ligt *et al*, 2012; Tavassoli *et al*, 2014; Fromer *et al*, 2014; Hoischen *et al*, 2014; Li *et al*, 2016; Johnson *et al*, 2016; Carroll *et al*, 2016; Balakrishna *et al*, 2020]. Consistent with these clinical findings, rodent models with *Scn2a* haploinsufficiency exhibit epileptic phenotypes, cognitive impairments, and behavioral abnormalities reminiscent of *SCN2A*-associated disorders in humans [Ogiwara *et al*., 2018; Middleton *et al*., 2018; Tatsukawa *et al*., 2019; Suzuki *et al*., 2024]. Our previous studies using genetic models of *SCN2A-* and *STXBP1*-related epilepsies demonstrated that reduced excitatory transmission from cortical pyramidal neurons onto striatal parvalbumin-expressing (PV^+^) fast-spiking interneurons (FSIs) is sufficient to trigger epilepsy [Ogiwara *et al*., 2018; Miyamoto *et al*., 2019]. Consistently, pharmacological inhibition of cortico-striatal excitatory inputs onto FSIs using Ca²⁺-permeable AMPA receptor antagonist induces generalized seizures in both rodents and non-human primates [Gittis *et al*., 2011; Aupy *et al*., 2024]. More recently, we further demonstrated that selective suppression of PV^+^ FSIs in the anteromedial shell of the NAc is sufficient to elicit convulsive seizures, underscoring the critical role of striatal microcircuits in seizure generation [Suzuki *et al*., 2026]. These findings highlight the importance of cortico-striatal circuits and PV⁺ FSIs in regulating network excitability. Accumulating evidence also suggests that *SCN2A* dysfunction contributes to behavioral abnormalities relevant to psychiatric disorders. For example, mice with *Scn2a* deficiency in brain regions implicated in schizophrenia and ASD, such as the medial prefrontal cortex (mPFC) and the VTA, display alterations in prepulse inhibition (PPI) of the acoustic startle response [Suzuki *et al*., 2024]. PPI is widely recognized as an endophenotype of schizophrenia [Braff *et al*., 2001; Geyer *et al*., 2006; Powell *et al*., 2009; Takahashi *et al*., 2011; Mena *et al*., 2016]. In addition, mice lacking *Scn2a* in the mPFC exhibit increased sociability, decreased locomotor activity, and enhanced anxiety-like behavior, whereas mice lacking *Scn2a* in the VTA show minimal behavioral abnormalities apart from altered vertical activity. These observations suggest that *Scn2a* dysfunction in distinct brain regions may differentially contribute to schizophrenia- and ASD-related phenotypes. In contrast, other group reported that deletion of *Scn2a* in VTA dopaminergic neurons reduces neuronal firing and dopamine release, leading to hyperactivity, impaired sociability, and reduced anxiety-like behavior [Li *et al*., 2025]. Similar abnormalities are observed in systemic *Scn2a* heterozygous knockout mice (*Scn2a*^+/-^) and are alleviated by acute levodopa administration, indicating dopamine system hypofunction [Li *et al*., 2025].

Based on these findings, dysfunction of *SCN2A*-dependent neural circuits, including cortico-striatal and mesolimbic dopamine pathways, has been proposed to contribute to the pathophysiology of ASD and schizophrenia. However, the circuit mechanisms through which *Scn2a* dysfunction leads to social behavioral abnormalities remain incompletely understood. In the present study, we sought to elucidate the neural circuit mechanisms underlying ASD-related social behavioral deficits associated with *SCN2A* dysfunction. In particular, we focused on the NAc, a key node in cortico-limbic circuits that integrates excitatory inputs from the cerebral cortex, hippocampus, and amygdala and regulates emotional and motivational behaviors. Using conditional genetic and chemogenetic approaches, we show that *Scn2a* haploinsufficiency in dorsal telencephalic excitatory neurons and selective inhibition of NAc PV⁺ FSIs reduce sociability. Furthermore, these behavioral abnormalities occur without detectable reductions in dopamine release in the NAc, suggesting that impaired activity of excitatory neurons or FSIs in cortico-striatal circuits may contribute to social behavioral deficits independently of mesolimbic dopamine hypofunction.

## Results

### Dorsal telencephalic excitatory neuron dysfunction and NAc FSI inhibition impair sociability

To selectively delete the *Scn2a* gene in excitatory neurons of the dorsal telencephalon, including the cerebral cortex, hippocampus, olfactory bulb, and amygdala, we generated conditional knockout mice (*Scn2a*^fl/+^/*Emx1*-Cre) using *Emx1*-Cre driver mice [Iwasato *et al*., 2000; Iwasato *et al*., 2004; Ogiwara *et al*., 2018], which mediate excitatory neuron-specific recombination in these regions (**Fig. 1A, B**). We then assessed sociability using the three-chamber social interaction test. Control mice showed a significant preference for the cage containing a stranger mouse over the empty cage. In contrast, mutant mice spent less time near the stranger mouse, indicating a reduced preference for the stranger mouse (**Fig. 1C**). These results indicate that *Scn2a* haploinsufficiency in dorsal telencephalic excitatory neurons impairs sociability and reduces interest in social stimuli.

**Figure 1.**
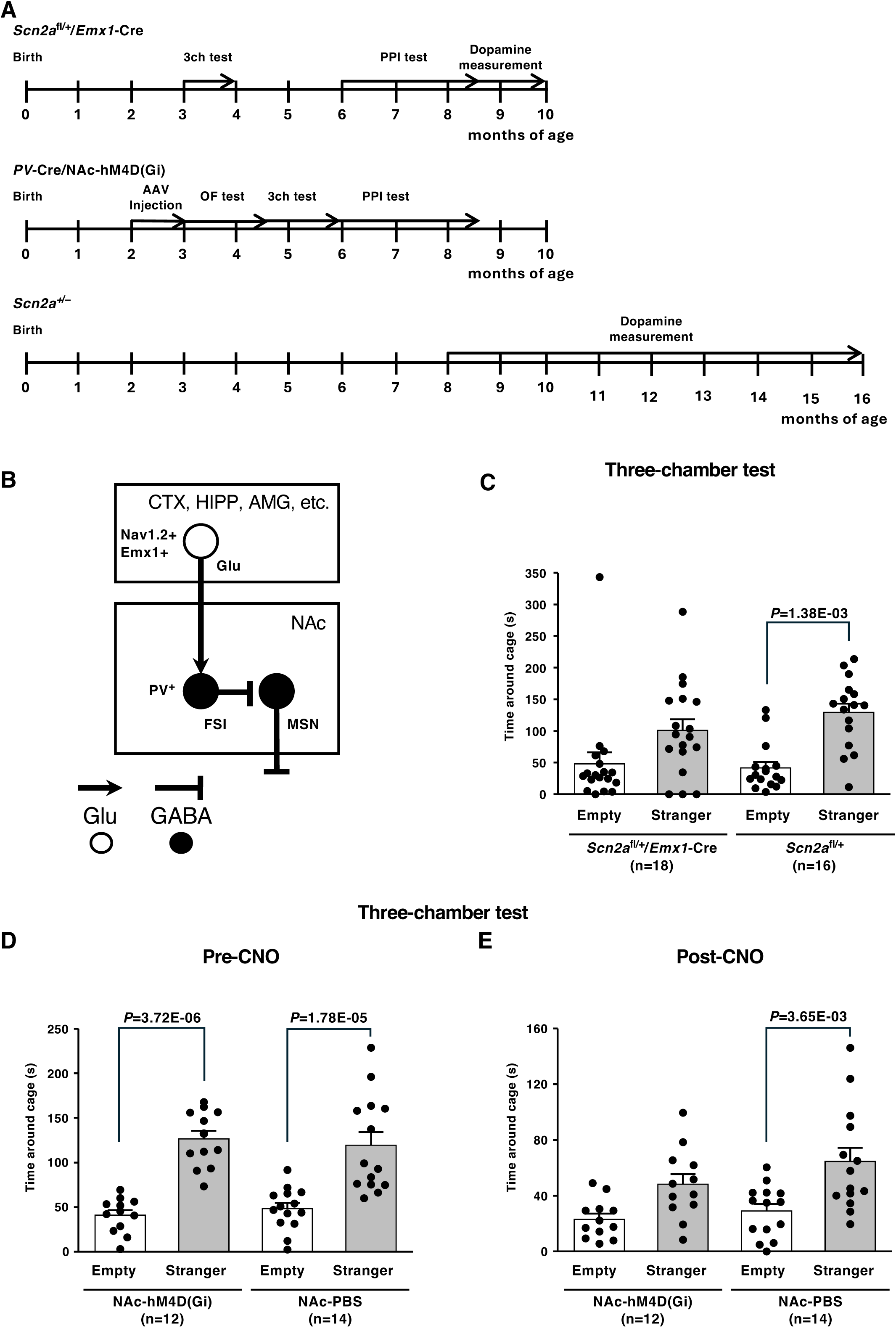
Reduced cortico-accumbal excitatory input and suppression of FSIs in the NAc impair sociability. (**A**) Schematic of the experimental design. (**B**) Schematic illustration of neural circuits projecting to the NAc. (**C**) During the sociability phase of the three-chamber social approach test, *Scn2a*^fl/+^/*Emx1*-Cre mice spent less time around the cage containing a stranger mouse, whereas control mice showed a clear preference for the cage containing the stranger mouse over the empty cage. (**D, E**) In the sociability phase of the three-chamber test, *PV*-Cre/NAc-hM4Di mice exhibited normal sociability before CNO administration (Pre-CNO; D), but showed reduced sociability after CNO administration (Post-CNO; E). Black dots represent individual mice. The number of mice in each group is indicated in parentheses. 3ch test, three-chamber test; AAV, adeno-associated virus; AMG, amygdala; CNO, clozapine-N-oxide; CTX, neocortex; FSI, fast-spiking interneuron; HIPP, hippocampus; MSN, medium spiny neuron; NAc, nucleus accumbens; OF test, open field test; PBS, phosphate-buffered saline; PPI test, prepulse inhibition test; PV^+^, parvalbumin-positive.

Next, we examined whether reduced activity of PV⁺ FSIs in the NAc produces a similar behavioral phenotype. The NAc receives projections from subsets of excitatory neurons in the cerebral cortex, hippocampus, and amygdala [Scofield *et al*., 2016; Marinescu *et al*., 2024; Xu *et al*, 2024] (**Fig. 1B**) and plays a key role in emotional and motivational processing [Castro et al., 2019]. To suppress NAc PV⁺ FSI activity, we injected an adeno-associated virus (AAV) encoding the inhibitory DREADD receptor hM4D(Gi) into the NAc of *PV*-Cre mice (*PV*-Cre/NAc-hM4Di). In this line, Cre recombinase is selectively expressed in PV⁺ FSIs, enabling targeted expression of hM4D(Gi). Histological analyses using the same mice confirmed that the AAV injections were accurately targeted to the intended region, as we previously reported [Suzuki *et al*., 2026]. We then performed the three-chamber test. Before clozapine-N-oxide (CNO) administration, *PV*-Cre/NAc-hM4Di mice displayed normal sociability, similar to control mice (**Fig. 1D**). After CNO administration, however, sociability was reduced, and mice spent less time near the cage containing the stranger mouse (**Fig. 1E**). Together, these results suggest that both reduced excitatory neuronal function in the dorsal telencephalon and decreased activity of NAc PV⁺ FSIs impair sociability.

### Selective *Scn2a* heterozygous deficiency in dorsal telencephalic excitatory neurons shows a trend toward reduced PPI

We next assessed PPI in *Scn2a*^fl/+^/*Emx1*-Cre mice. Startle responses to acoustic stimuli at two intensities (110- and 120-dB) were comparable between mutant and control (*Scn2a*^fl/+^) mice (**Fig. 2A**). In contrast, the percentage PPI of the startle response to a 120-dB pulse with 74- and 78-dB prepulses tended to be lower in mutant mice, although the differences did not reach statistical significance (**Fig. 2B**).

**Figure 2.**
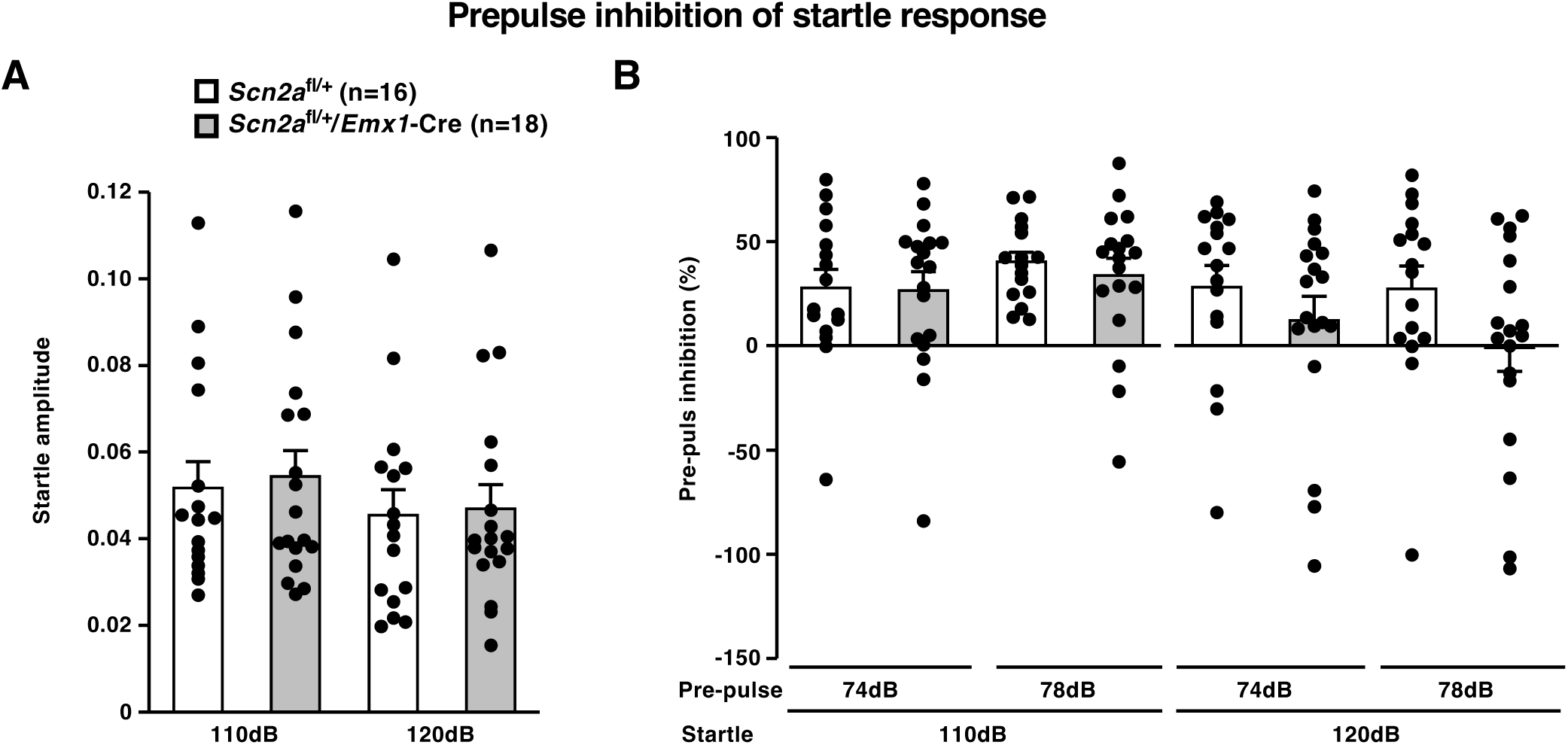
Selective *Scn2a* deficiency in dorsal telencephalic excitatory neurons shows a trend toward reduced PPI. (**A**) Acoustic startle responses to two sound stimulus intensities (110- and 120-dB) did not differ significantly between *Scn2a*^fl/+^/*Emx1*-Cre mice and control *Scn2a*^fl/+^ mice. (**B**) In the PPI test, the percentage PPI of the startle response to a 120-dB pulse preceded by 74- or 78-dB prepulses showed a trend toward reduction in *Scn2a*^fl/+^/*Emx1*-Cre mice compared with control mice. Black dots represent individual mice, and the number of mice is indicated in parentheses.

### Inactivation of NAc FSIs has no effect on locomotor activity or anxiety-like behavior

We next assessed spontaneous exploratory behavior using the open field test. *PV*-Cre/NAc-hM4Di mice showed no significant differences in total distance traveled (**Fig. 3A**) or time spent in the center zone (**Fig. 3B**), either before or after CNO administration. These results indicate that chemogenetic inactivation of NAc FSIs does not affect locomotor activity or anxiety-like behavior under these conditions.

**Figure 3.**
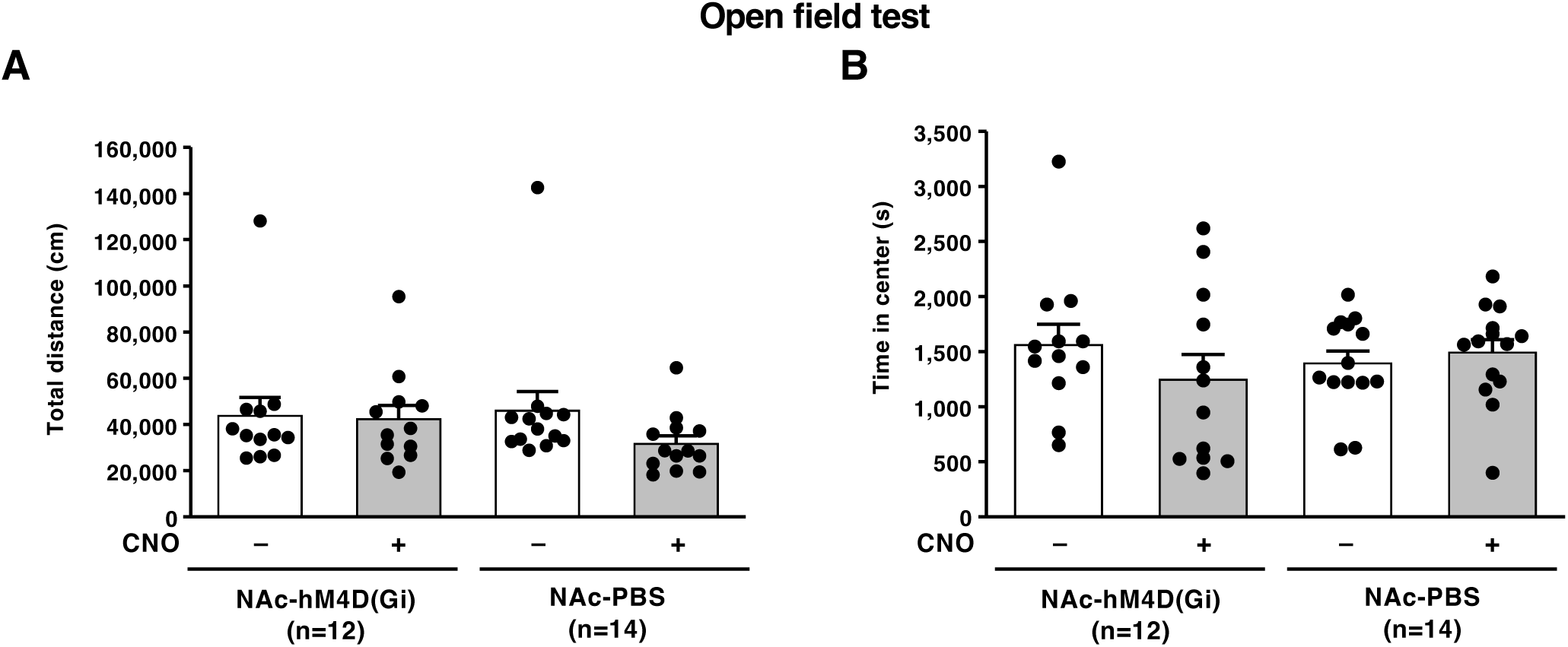
Inactivation of NAc FSIs does not affect exploratory or anxiety-like behavior in the open field test. (**A**) Total distance traveled in the open field test did not differ between *PV*-Cre/hM4Di mice and control mice injected with PBS into the NAc, either before (−) and after (+) CNO administration. (**B**) Time spent in the center zone of the open field area was comparable between *PV*-Cre/hM4Di mice and control mice injected with PBS into the NAc, both before (−) and after (+) CNO administration. Black dots represent individual mice, and the number of mice in each group is indicated in parentheses.

### Dopamine release in the NAc is comparable between *Scn2a* mutant and control mice

Finally, we measured dopamine (DA) release in the NAc under novel environmental conditions and following methamphetamine administration. Although our experimental procedures differed from those of a previous study [Li *et al*., 2025], DA release in *Scn2a*^fl/+^/*Emx1*-Cre mice (**Fig. 4A, B**) and *Scn2a*^+/-^ mice (**Fig. 4C, D**) was comparable to that in control mice. These findings suggest that the sociability deficits observed in *Scn2a*^fl/+^/*Emx1*-Cre mice and *PV*-Cre/NAc-hM4Di mice cannot be explained solely by reduced DA release in the NAc. Instead, they raise the possibility that altered activity in other circuits, such as dorsal telencephalic excitatory neurons or NAc FSIs, also contributes to social behavioral deficits.

**Figure 4.**
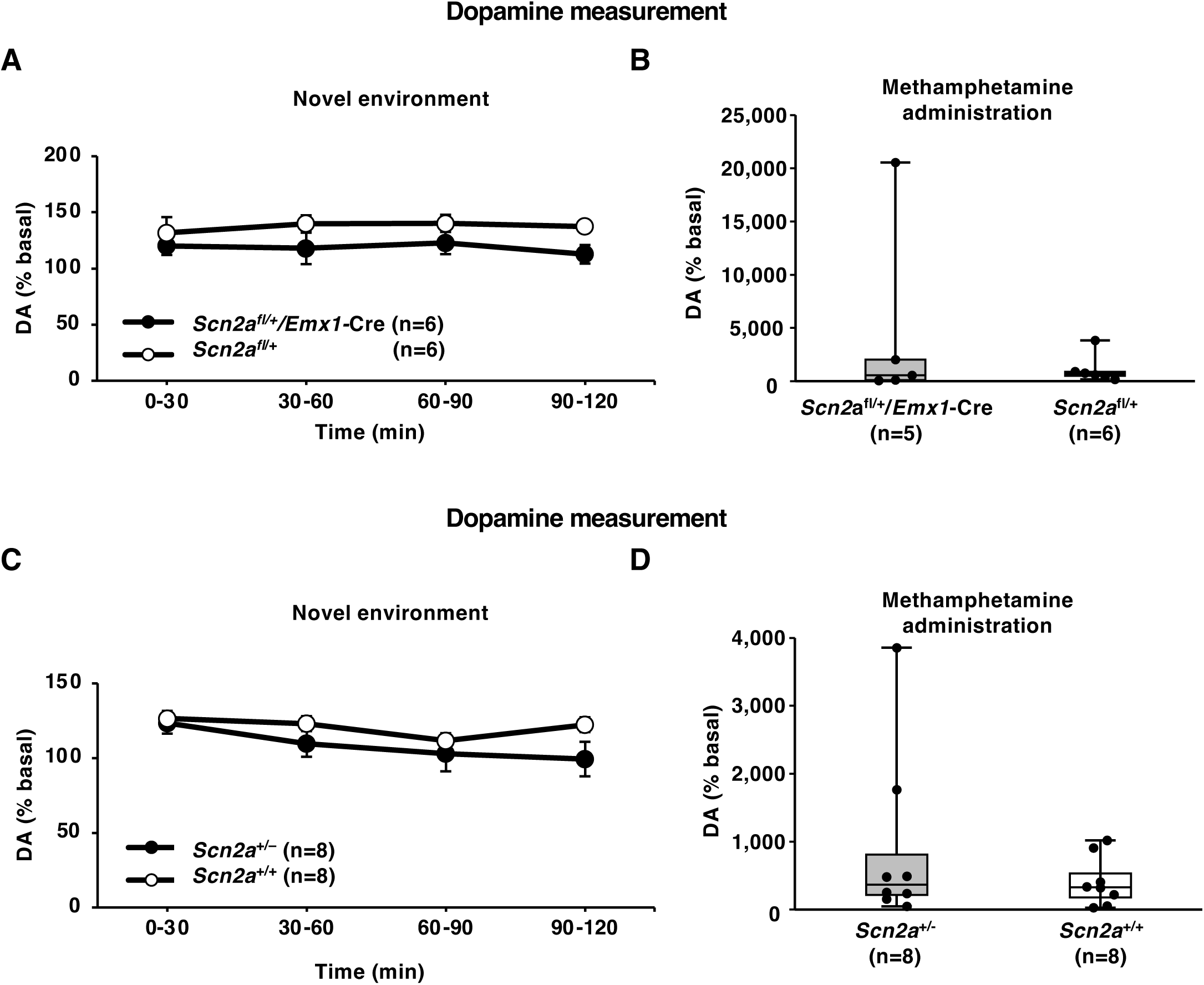
Dopamine release in the NAc is comparable between *Scn2a* mutant and control mice. Dopamine release in the NAc of *Scn2a*^fl/+^/*Emx1*-Cre mice (**A, B**) and *Scn2a*^+/-^ mice (**C, D**) was measured under a novel environmental condition (A, C) and following methamphetamine administration (B, D). Data are presented as mean ± s.e.m. (A, C) or as box-and-whisker plots (B, D). Black dots represent individual mice (B, D), and the number of mice in each group is indicated in parentheses (A–D). DA, dopamine.

## Discussion

The present study provides new insight into the circuit mechanisms underlying social behavioral deficits associated with *Scn2a* dysfunction. Using conditional genetic and chemogenetic approaches, we show that reduced activity of dorsal telencephalic excitatory neurons or NAc PV⁺ FSIs is sufficient to impair sociability. Notably, these behavioral alterations occur in the absence of detectable changes in DA release in the NAc, indicating that dysfunction of cortico-accumbal microcircuits, rather than mesolimbic DA hypofunction alone, can drive social deficits.

A key finding is that *Scn2a* haploinsufficiency in dorsal telencephalic excitatory neurons reduces sociability, in contrast to previous reports of increased social interaction in mice with systemic *Scn2a* haploinsufficiency or mPFC-specific deletion [Tatsukawa *et al*., 2019; Suzuki *et al*., 2024; Li *et al*., 2025]. This discrepancy underscores the circuit-dependent roles of *Scn2a* in social behavior. One critical difference lies in the extent and composition of affected circuits. Whereas mPFC-specific deletion primarily alters prefrontal outputs, *Emx1*-Cre–mediated deletion targets a broader population of excitatory neurons across the cortex, hippocampus, and amygdala. These regions provide convergent glutamatergic inputs to the NAc, a central hub for integrating social and motivational signals. Thus, a widespread reduction in excitatory drive may decrease the salience or reward value of social stimuli, leading to reduced sociability. Consistent with this interpretation, direct chemogenetic inhibition of NAc PV⁺ FSIs recapitulated the social deficit phenotype. PV⁺ FSIs provide strong feedforward inhibition onto MSNs and regulating the timing and gain of striatal output. Reduced FSI activity is therefore likely to disrupt the excitation-inhibition balance within the NAc, impairing the encoding of socially relevant information. Together, these findings support model in which *Scn2a* dysfunction produces bidirectional effects on sociability depending on the affected circuit, with cortico-accumbal hypofunction leading to social deficits, in contrast to hyper-sociability observed following more restricted perturbations.

In addition to social behavior, *Scn2a* haploinsufficiency in dorsal telencephalic excitatory neurons showed a trend toward reduced PPI. This is consistent with our previous finding that mPFC-specific *Scn2a* deletion impairs PPI [Suzuki *et al*., 2024], supporting a central role for cortical excitatory circuits in sensorimotor gating. Given that the *Emx1*-Cre driver includes mPFC neurons, the observed PPI deficit likely reflects impaired prefrontal function. The convergence of PPI deficits across dorsal telencephalic and mPFC-specific manipulations further suggests that distributed cortical network dysfunction, rather than a single locus, underlies impaired sensorimotor gating. In contrast, the enhancement of PPI observed following *Scn2a* deletion in VTA dopaminergic neurons [Suzuki *et al*., 2024] highlights the distinct and potentially opposing contributions of cortical and dopaminergic systems.

Chemogenetic inhibition of NAc PV⁺ FSIs did not significantly alter locomotor activity or anxiety-like behavior in the open field test. This contrasts with previous reports showing hyperactivity in systemic *Scn2a* haploinsufficiency [Tatsukawa *et al*., 2019] and reduced locomotion following mPFC-specific deletion [Suzuki *et al*., 2024]. These differences are consistent with the notion that the NAc is more closely associated with motivational and emotional processing than with general motor control. Accordingly, PV⁺ FSIs in the NAc may preferentially regulate socially and motivationally relevant behaviors. Alternatively, compensatory mechanisms within striatal circuits may preserve locomotor function, or subtle behavioral changes may not be captured by the open field paradigm.

Based on these findings, we propose a circuit model (**Fig. 5**) in which reduced excitatory drive from the dorsal telencephalon leads to decreased activation of PV⁺ FSIs, resulting in dysregulated striatal output. Because FSIs receive dense glutamatergic inputs from cortical and limbic regions, reduced excitatory neuronal activity is likely to indirectly suppress FSI function. This, in turn, may alter activity patterns in MSNs and disrupt downstream signaling within ventral striatal circuits. Such disruption could impair neural processes underlying reward prediction, social salience, and action selection. Notably, our previous work demonstrated that impaired cortico-striatal excitation onto the striatal FSIs contributes to epileptogenesis [Miyamoto *et al*., 2019], suggesting that PV⁺ FSIs represent a critical convergence point linking network excitability and behavioral regulation. Thus, NAc PV⁺ FSIs may function as an integrative hub through which cortical deficits are translated into behavioral abnormalities, including both social deficits and impaired PPI.

**Figure 5.**
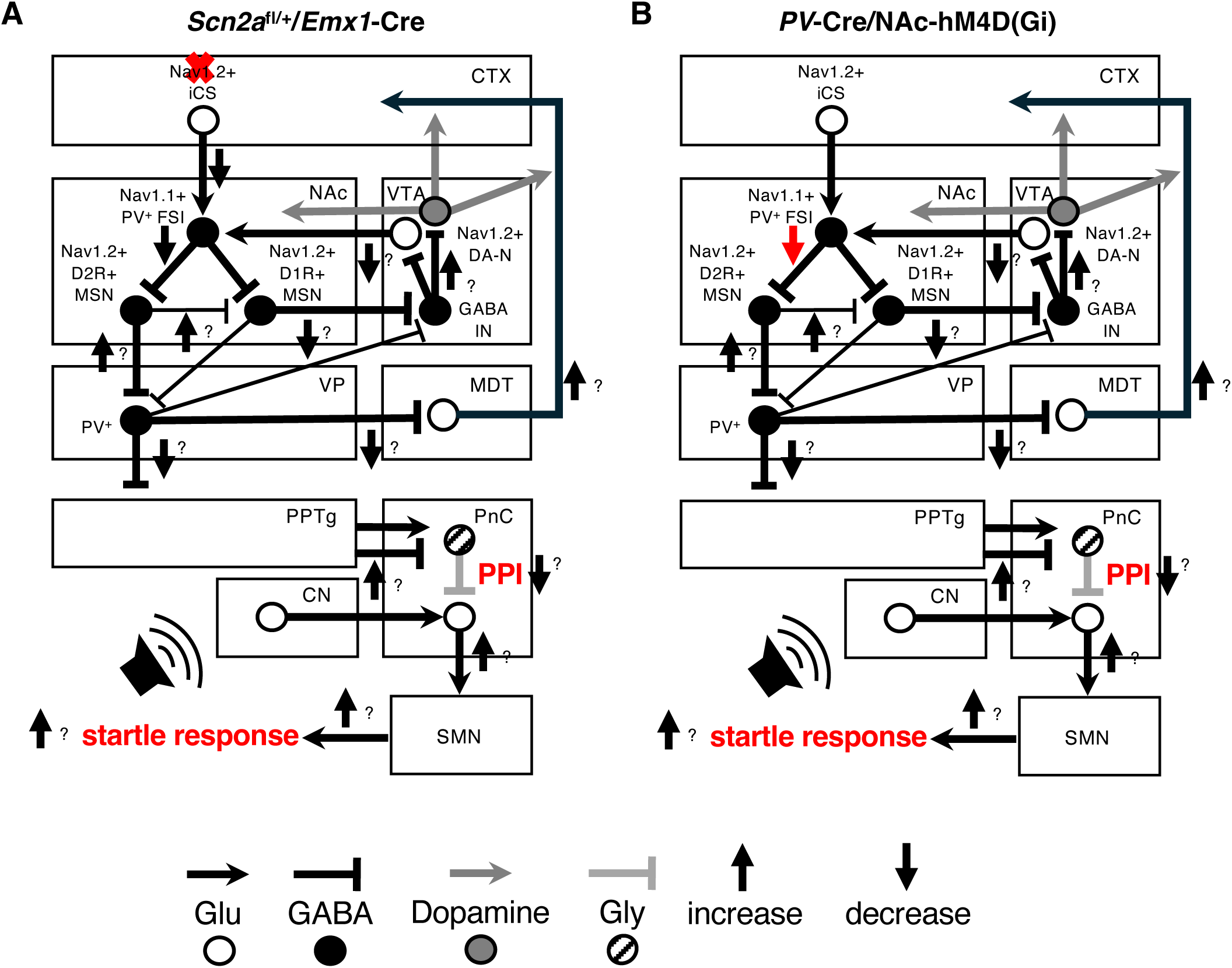
Cortico-accumbal circuit models for sociability and PPI in *Scn2a* mutant mice. Schematic models of neural circuit underlying sociability and prepulse inhibition (PPI) in *Scn2a* mutant mice. These diagrams are modified and simplified from previous published models [Swerdlow *et al*., 2001; Miyamoto *et al*., 2019; Cano *et al*., 2021; Suzuki *et al*., 2024; Suzuki *et al*., 2026], and incorporate interpretations based on the present findings. Hypothetical neural circuit models illustrating the mechanisms underlying reduced PPI in *Scn2a*^fl/+^/*Emx1*-Cre mice (**A**) and *PV*-Cre/hM4Di mice (**B**). Upward and downward arrows indicate increased and decreased neural transmission, respectively. CN, cochlear nuclei; CTX, neocortex; D1R, dopamine D1 receptor; D2R, dopamine D2 receptor; DA-N, dopaminergic neuron (gray circle) and transmission (gray arrows); FSI, fast-spiking interneuron; GABA, GABAergic neurons (black circles) and transmission (black lines); Glu, glutamatergic neurons (white circles) and transmission (black arrows); Gly, glycinergic neurons (striped circle) and transmission (gray lines); iCS, intratelencephalic cortico-striatal neurons; IN, interneurons; MDT, mediodorsal thalamus; MSN, medium spiny neuron; NAc, nucleus accumbens; Nav1.1, voltage-gated sodium channel α1 subunit; Nav1.2, voltage-gated sodium channel α2 subunit; PnC, caudal pontine reticular nucleus; PPI, prepulse inhibition; PPTg, pedunculopontine tegmental nucleus; SMN, spinal motor nerve; VP, ventral pallidum; VTA, ventral tegmental area.

Recent work by Li *et al*. (2025) demonstrated that *Scn2a* deletion in VTA dopaminergic neurons reduces DA release in the NAc and leads to social deficits, supporting a DA-dependent mechanism. In contrast, we found that DA release in the NAc is largely preserved in both dorsal telencephalon-specific and systemic *Scn2a* mutant mice despite impaired sociability. These findings indicate that social deficits in *Scn2a* models cannot be explained solely by mesolimbic DA hypofunction. Instead, they support the existence of at least two parallel pathways: (1) a DA-dependent pathway involving VTA dysfunction, and (2) a DA-independent pathway involving cortico-striatal circuit dysfunction. These mechanisms are not mutually exclusive and may interact. For example, cortical inputs could modulate DA neuron activity indirectly, while DA signaling may influence striatal microcircuit dynamics. In addition, the VTA provides glutamatergic projections to the NAc, which may also contribute to motivated behavior [Qi *et al*., 2016]. Although not examined here, *Scn2a* dysfunction in these projections could represent an additional mechanism and warrants further investigations.

In summary, our findings demonstrate that reduced activity of dorsal telencephalic excitatory neurons or NAc PV⁺ FSIs is sufficient to impair sociability, independently of changes in NAc dopamine release. These results highlight the importance of cortico-accumbal microcircuits in social behavior and support a framework in which *SCN2A*-related neuropsychiatric phenotypes arise from circuit-specific dysfunction across distributed brain networks. Targeting striatal microcircuits or restoring excitation-inhibition balance may therefore represent promising therapeutic strategies for *SCN2A*-assiciated disorders.

## Methods

### Animals

All mice were maintained under a 12 h light/dark cycle with ad libitum access to food and water. *Scn2a* floxed mice [Ogiwara *et al*., 2018], in which exon 2 is flanked by loxP sites, and Empty spiracles homolog 1 (Emx1)-Cre knock-in mice [Iwasato *et al*., 2000; Iwasato *et al*., 2004] were maintained on a C57BL/6J background. To generate *Scn2a* conditional knockout mice, homozygous *Scn2a* floxed (*Scn2a*^fl/fl^) mice were crossed with *Emx1*-Cre knock-in mice. Parvalbumin (PV)-Cre transgenic mice [Tanahira *et al*., 2009] were maintained on a C57BL/6J background. *Scn2a* heterozygous knock-out (*Scn2a*^+/-^) mice were described previously [Planells-Cases *et al*., 2000] and were maintained on a C57BL/6J background.

### AAVs

The plasmid construct for double floxed Gi-coupled hM4D DREADD fused with mCherry under the control of EF1a promoter (pAAV-EF1a-DIO-hM4D(Gi)-mCherry) was a gift from Dr. Bryan Roth (Addgene plasmid 50461). Packaging of the plasmid vector into AAVs, as well as purification and quantification of the viruses, were performed at the Division of Genetic Therapeutics, Jichi Medical University, and at the section for Viral Vector Development, National Institute of Physiological Sciences.

### Stereotaxic surgery

*PV*-Cre transgenic mice (>8 weeks of age, both sexes) were anesthetized with isoflurane (1.0–2.5%) and placed in a stereotaxic apparatus (Stoelting). Bilateral injections of AAV5-EF1α-DIO-hM4D(Gi)-mCherry (7.5 × 10^12^ viral genomes/ml; 250 nl per site) or phosphate-buffered saline (PBS; 250 nl per site) into the NAc were performed using a microinjector (Nanoliter 2020 Injector; World Precision Instruments) fitted with a pulled glass capillary at a flow rate of 100 nl/min. Stereotaxic coordinates were determined based on a mouse brain atlas [Paxinos *et al*., 2001]: NAc (anteroposterior, mediolateral, and dorsoventral, in mm): +1.60, ±1.00, −5.00 and −4.50; and +0.80, ±1.00, −5.20 and −4.50.

### Measurements of dopamine release

Extracellular dopamine levels in the NAc of male mice aged 8–15 months were quantified utilizing an in vivo microdialysis system. Under anesthesia induced by a combination of medetomidine (0.3 mg/kg), midazolam (4 mg/kg), and butorphanol (5 mg/kg), a guide cannula (AG-6; EICOM) was implanted into the NAc shell. The coordinates for implantation, based on the mouse brain atlas [Paxinos *et al*., 2001], were as follows (in mm): anteroposterior +1.30, mediolateral +0.70, dorsoventral −4.20. The cannula was affixed to the skull using stainless steel screws and dental acrylic cement. Three to four days post-surgery, a dialysis probe equipped with a 1 mm-length membrane (FX-I-6-01, EICOM) was inserted into the guide cannula. This probe was perfused with artificial cerebrospinal fluid, composed of 147.2 mM NaCl, 4.0 mM KCl, and 2.3 mM CaCl_2_, at a flow rate of 0.8 μl/min using a microsyringe injector (ESP-64, EICOM). Dialysate samples were collected and injected every 30 minutes using an auto injector (EAS-20S, EICOM). The samples were subsequently separated using an SC-50DS column (EICOM) to facilitate dopamine measurement via high-performance liquid chromatography (HPLC) with an electrochemical detection system (HTEC-500, EICOM), where the working electrode was maintained at +700 mV (vs. Ag/AgCl) at a temperature of 25 °C. The mobile phase consisted of 83 % 0.1 M citrate/0.1 M sodium acetate buffer (pH 3.9) and 17 % methanol, containing 140 mg/l sodium 1-octanesulfonate and 5 mg/l EDTA-2Na, delivered at a flow rate of 0.23 ml/min. Protocol was controlled and data analysis was performed using Power Chrom software (ADI instruments Japan).

The animal was acclimated for over two hours in the experimental arena, which measured 30 × 30 × 35 cm, until the baseline stabilized. Following the two-hour measurement period (four samples), the animals were administered methamphetamine (1.0 mg/kg, i.p.), and samples were obtained for more than two hours.

### Three-chamber social approach test

The three-chamber apparatus consisted of a rectangular clear Plexiglas box (43 × 63 cm) divided into three equal-sized chambers (21 × 43 cm) by transparent Plexiglas partitions with small openings that allowed mice to freely move between chambers. A cylindrical wire cage was placed in each corner of the two side chambers to enclose an unfamiliar 11–12-week-old C57BL/6J male mouse (stranger). Test male mice (3–5 months of age) were first habituated to the apparatus for 10 min, with an empty wire cage placed in each of the side chambers. For the sociability test, a wire cage containing a stranger mouse was placed in one side chamber, and the subject mouse was initially placed in the center chamber and allowed to freely explore the entire apparatus for a 10 min. The location of the stranger mouse (left or right chamber) was alternated between trials. Behavior was recorded using a video camera mounted above the apparatus. The total time spent in proximity to each cage (empty or containing the stranger mouse) was quantified manually in a blinded manner.

### Startle response and prepulse inhibition (PPI) test

The startle response and PPI test was conducted as previously described [Nakao *et al*., 2015; Suzuki *et al*., 2024]. Male mice (6–8 months of age) were placed in a Plexiglas cylinder with a continuous 70 dB white noise background and allowed to habituate for 5 min. An acoustic startle stimulus (40 ms; 110- or 120-dB) was presented either alone or preceded by a prepulse stimulus (20 ms; 74- or 78-dB) during the PPI trials. Each test block consisted of two startle alone trials (110- and 120-dB) and four prepulse–startle trials (74-dB prepulse + 110- or 120-dB startle; 78-dB prepulse + startle 110- or 120-dB startle). Each mouse underwent ten blocks. The inter-trial interval varied randomly with an average of 15 s (range: 10–20 s). Startle responses and PPI were recorded automatically.

### Open field test

Male mice (3–4 months of age) were placed in a corner of a square open-field apparatus (45 × 45 × 20 cm) illuminated at 100 lx and allowed to freely explore for 120 min. The total distance traveled (cm) and the time spent in the center area (20 × 20 cm) were automatically recorded using manufacturer’s software (Smart3.0, PanLab).

### Statistical analyses

Data were analyzed using Student’s t-test for two-group comparisons or one-way or two-way analysis of variance (ANOVA), followed by Tukey’s post hoc test for parametric data (KyPlot; KyensLab Inc.), as appropriate. Data are presented as the mean ± standard error of the mean (s.e.m.) or as box-and-whisker plots. In the box-and- whisker plots, the boxes indicate the median (thick line), the 25th and 75th percentiles, and the whiskers represent the minimum and maximum values. Statistical significance was defined as P < 0.05.

## Acknowledgments

The authors thank Drs. Tamamaki and Tanahira (Kumamoto University) for generously providing the PV-Cre-TG driver line. We also thank the members of the Department of Neurophysiology and Brain Science, the Endowed Department of Cognitive Function and Pathology, the Department of Neurodevelopmental Disorder Genetics, and the staff of the animal facility at Nagoya City University (NCU) for their support. We are grateful to the Research Equipment Sharing Center at NCU for technical assistance.

## Author Contributions

TS and KaY contributed to the study conception and design. Material preparation, data collection, and analysis were performed by TS, ST, YY, HM, KK, WN, KoY, TK, YH, TY, SI, HN, HH, KaY. The first draft of the manuscript was written by TS and KaY. All authors have reviewed and approved the final manuscript.

## Funding

This work was supported by grants from NCU; JSPS KAKENHI (Grant Numbers 22K07620 to TS, 23K27490 to KaY and 25K10820 to TS); and the Grant-in-Aid for Outstanding Research Group Support Program at NCU (Grant Number 2401101). This study also utilized research equipment shared under the MEXT Project for Promoting Public Utilization of Advanced Research Infrastructure (Program for Supporting Construction of Core Facilities; Grant Number JPMXS0441500024).

## Data Availability

The datasets generated during and/or analyzed during the current study are available from the corresponding author on reasonable request.

## Declarations Ethics Approval

All animal breeding and experimental procedures were approved by the Institutional Animal Care and Use Committee of NCU (approval No. 19-032, approved 20 Dec 2022; approval No. 23-024, approved 21 Apr 2023). All procedures were conducted in accordance with the ARRIVE guidelines and the institutional guidelines and regulations of NCU.

## Consent to Participate

Not applicable.

## Consent for Publication

Not applicable.

## Competing Interests

The authors declare no competing interests.

## Notes

### Competing Interest Statement

The authors have declared no competing interest.

